# Communication between the mediodorsal thalamus and prelimbic cortex regulates timing performance in rats

**DOI:** 10.1101/2021.06.18.449036

**Authors:** Benjamin J. De Corte, Kelsey A. Heslin, Nathan S. Cremers, John H. Freeman, Krystal L. Parker

**Author notes:** **Corresponding Author**, Department of Psychiatry, University of Iowa, Iowa City, IA, USA.

## Abstract

Predicting when future events will occur and adjusting behavior accordingly is critical to adaptive behavior. Despite this, little is known about the brain networks that encode time and how this ultimately impacts decision-making. One established finding is that the prefrontal cortex (PFC) and its non-human analogues (e.g., the rodent prelimbic cortex; PL) mediate timing. This provides a starting point for exploring the networks that support temporal processing by identifying areas that interact with the PFC during timing tasks. For example, substantial work has explored the role of frontostriatal circuits in timing. However, other areas are undoubtedly involved. The mediodorsal nucleus of the thalamus (MD) is an excellent candidate region. It shares dense, reciprocal connections with PFC-areas in both humans and non-human species and is implicated in cognition. However, causal data implicating MD-PFC interactions in cognition broadly is still sparse, and their role in timing specifically is currently unknown. To address this, we trained male rats on a time-based, decision-making task referred to as the ‘peak-interval’ procedure. During the task, presentation of a cue instructed the rats to respond after a specific interval of time elapsed (e.g., tone-8 seconds). We incorporated two cues; each requiring a response after a distinct time-interval (e.g., tone-8 seconds / light-16 seconds). We tested the effects of either reversibly inactivating the MD or PL individually or functionally disconnecting them on performance. All manipulations caused a comparable timing deficit. Specifically, responses showed little organization in time, as if primarily guided by motivational systems. These data expand our understanding of the networks that support timing and suggest that MD-PL interactions specifically are a core component. More broadly, our results suggest that timing tasks provide a reliable assay for characterizing the role of MD-PL interactions in cognition using rodents, which has been difficult to establish in the past.

## Introduction

Time is fundamental to cognitive function. When making a decision, we not only select an appropriate action but also execute it at an appropriate time (Fan et al., 2012). When forming a new memory, we not only encode what happened and where, but also when the event occurred (Tulving, 1986). Furthermore, to comprehend causality, we must be able to recognize that causes precede their effects (Hume, 1739). Given its broad relevance, how individuals perceive time is discussed across many domains, such as philosophy (Mctaggart, 1908), psychology (Fraisse, 1964), neuroscience (Paton & Buonomano, 2018), and treating cognitive deficits in disease (Parker, 2015). However, despite its importance, we know little about the brain networks that encode time.

Part of the problem is that timing appears to rely on a diverse set of brain areas. For example, timing has been studied extensively in association cortices, the basal ganglia, and midbrain dopamine centers (Jazayeri & Shadlen, 2015; Mello et al., 2015; Soares et al., 2016). However, many other areas have been implicated such as the hippocampus (Shikano et al., 2021), cerebellum (Narain et al., 2018), and even the primary visual cortex (Shuler, 2006). Explaining how these areas work together to mediate timing is a difficult endeavor. A further complication is the diversity of time-related signals seen among neurons in most of these areas while individuals track time. For example, ‘ramping’ neurons that monotonically change in firing rate as a learned time-interval elapses have been well documented (Chen et al., 2017; Jazayeri & Shadlen, 2015; Leon & Shadlen, 2003). However, ‘time-cells’ that show consistent Gaussian-like firing-profiles around specific time intervals have also been shown (Mau et al., 2018; Mello et al., 2015; Zhou et al., 2020). Furthermore, there is even indication that neurons encode time via idiosyncratic, non-linear firing profiles (Buonomano & Maass, 2009; Buonomano & Mauk, 1994; Wang et al., 2018).

Importantly, converging evidence suggests that the prefrontal cortex (PFC) is necessary for timing intervals in the seconds-to-minutes range—referred to as ‘interval timing’ (Matell & Meck, 2000). The PFC’s precise role in interval timing is still unclear. However, disruption to the human PFC or its non-human analogues causes disruption in a variety of timing tasks (Buhusi et al., 2018; Koch et al., 2003; Wang et al., 2018). Furthermore, all forms of time-related signaling noted above have been observed in this area (Matell & Meck, 2004; Wang et al., 2018).

Therefore, a useful approach for understanding the networks that support timing is to assess what areas interact with the PFC during timing tasks. For example, substantial focus has been dedicated to defining the role of frontostriatal circuits in timing (Buhusi & Meck, 2005; Matell & Meck, 2004). Broadly, the data suggests that the PFC encodes the passage of time, and the striatum uses this information to appropriately execute decisions (Matell et al., 2003; Mello et al., 2015). However, more recent data suggests that the frontal cortex and thalamus functionally interact during timing tasks (Parker et al., 2017; Wang et al., 2018). Nonetheless, which thalamic nuclei engage with the PFC during timing tasks has yet to be established.

One promising candidate region is the mediodorsal nucleus (MD) of the thalamus. Neuroanatomically, the MD shares dense, reciprocal projections with the PFC across species (Alcaraz et al., 2016; Georgescu et al., 2020; Ray & Price, 1993). Functionally, the MD plays a role in cognitive functions (Markowitsch, 1982; Peräkylä et al., 2017), although finding a robust, cross-species behavioral assay to test its role in cognition has been difficult (Brito et al., 1982; Mitchell & Chakraborty, 2013). Furthermore, the MD’s role in cognition appears to be mediated, in part, by the projections it shares with prefrontal areas (Giraldo-Chica et al., 2018; Pergola et al., 2013). Importantly, while data are limited at this point, some studies have shown that the MD itself is necessary for timing (Lusk et al., 2020; Yu et al., 2010).

Based on this, we developed the current study around the following goals. First, we sought to further establish the roles of the PL and MD in timing. Second, we sought causal evidence that MD-PL interactions mediate timing. To this end, we trained rats on a time-based decision-making task, wherein cues prompted subjects to respond after certain time-intervals elapsed. We incorporated two cues; each associated with a distinct time-interval (e.g., tone-8 seconds / light-16 seconds). We then inactivated the MD or PL individually or blocked MD-PL communication. Incorporating two time-intervals allowed us to evaluate the consistency of effects across durations. Furthermore, incorporating an auditory and visual cue allowed us to evaluate whether these areas play a sensory-specific or amodal role in timing. Along with the use of within-subjects designs, we were able to obtain a detailed assessment of how the MD, PL, and their interaction impact timing.

## Results

### The MD and PL are necessary for timing performance

We first assessed the necessity of the mediodorsal nucleus of the thalamus (MD) and prelimbic cortex (PL) to timing performance. Specifically, we trained nine rats on a timing task called the ‘peak-interval’ procedure (Fig 1A). During this task, trials start with the presentation of a cue that signals reward can be earned for responding after a specific interval of time has elapsed (e.g., tone-8s). During probe trials, no reward is delivered, and rats’ responses typically cluster around the entrained interval. In this experiment, we trained rats to associate two cues with distinct time intervals (e.g., tone-8s / light-16s), presenting each cue individually during separate trials. Once trained, we implanted cannulae bilaterally into the MD and PL (Fig 1C). We then tested the effects of reversibly inactivating either area during separate sessions using muscimol (Fig 1B).

**Figure 1.**
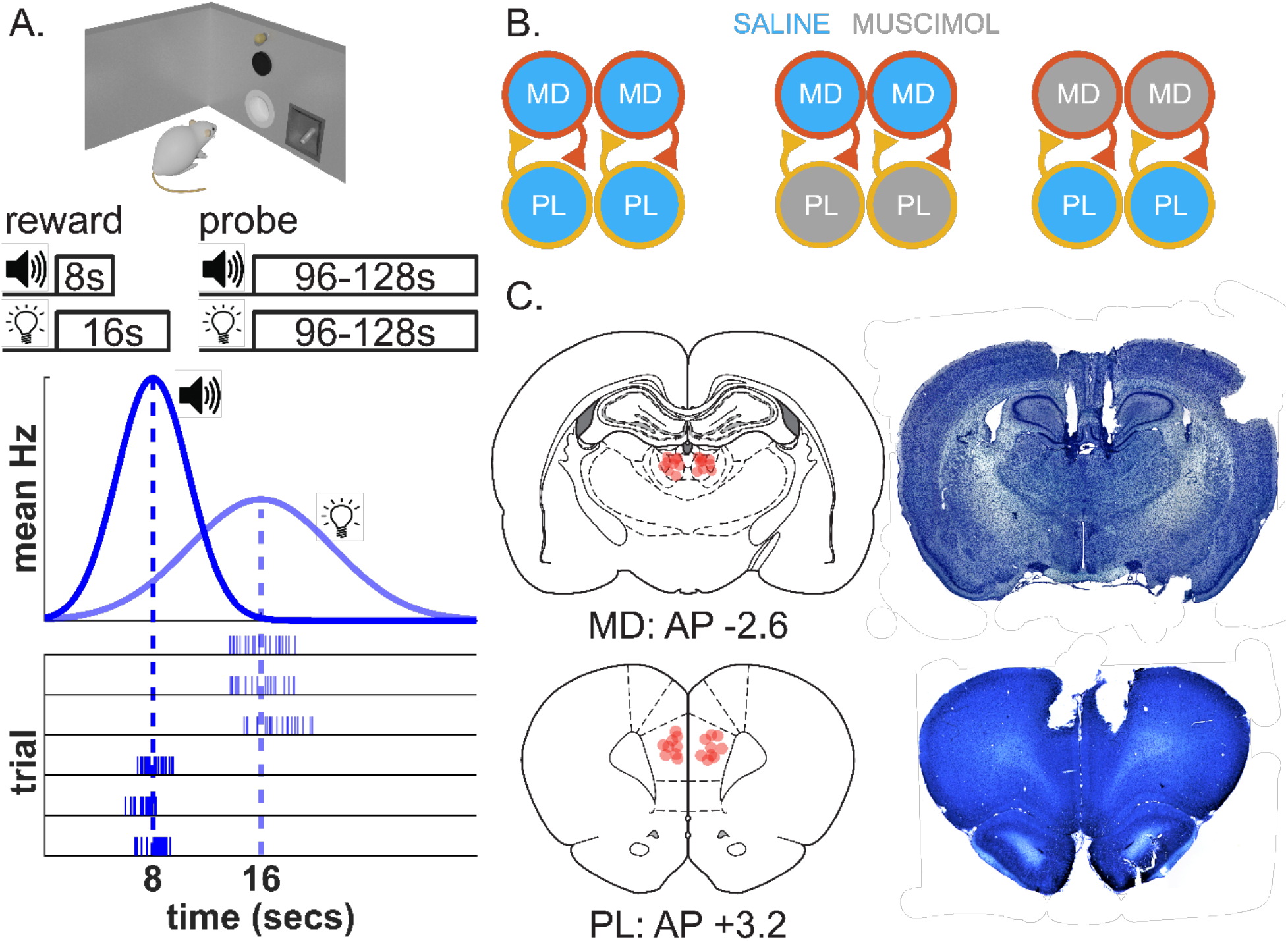
Task summary and histological analysis for within-subject bilateral inactivations of the MD and PL. (A) Description of task including a chamber schematic (houselight, tone generator, nosepoke, and reward port) and trial structure, illustrating rewarded and probe trials for both tone and light cues. Bottom panels illustrate the difference between patterns of responding in averaged response rate (gaussian curves, mean Hz) and single-trial behavior (bursts). (B) Infusion design for saline sessions, PL inactivations, and MD inactivations (left, middle, right, respectively). (C) Diagram of cannula tip placements in MD and PL, with representative histological images.

After saline infusions, mean response rates across probe trials showed typical, Gaussian-like curves, peaking near each cue’s respective interval (Fig 2A). However, inactivating the MD or PL with bilateral muscimol infusions flattened these curves, with responses being less common around the entrained intervals and more common at irrelevant times throughout the trial (Fig 2A). Specifically, the fit-quality of a Gaussian distribution to response curves dropped following MD and PL inactivations [Fig 2B; Infusion: *F*(2,14) = 15.51, *p* < 0.001; MD: *t*(6) = -4.50, *p* < 0.005; PL: *t*(6) = -4.79, *p* < 0.005; *R*^2^ *M* +/*SEM*: Sal = 0.90 +/-.03, MD = 0.48 +/-.08, PL = 0.74 +/-.03]. Furthermore, inactivations did not reliably change overall mean response rates during probe trials [Fig 2B; Infusion: *F*(2,14) = 0.24, *p* = 0.794]. Together, these data suggest that rats were unable to time their responses appropriately following MD and PL inactivations, with no evidence for decreased motivation or ability to respond, per se. Given the drop in fit-quality, the validity of measures derived from the fits is questionable (e.g., peak-time, spread). Nonetheless, for exploratory purposes, we provide standard analyses in the supplemental results (Fig S1).

**Figure 2.**
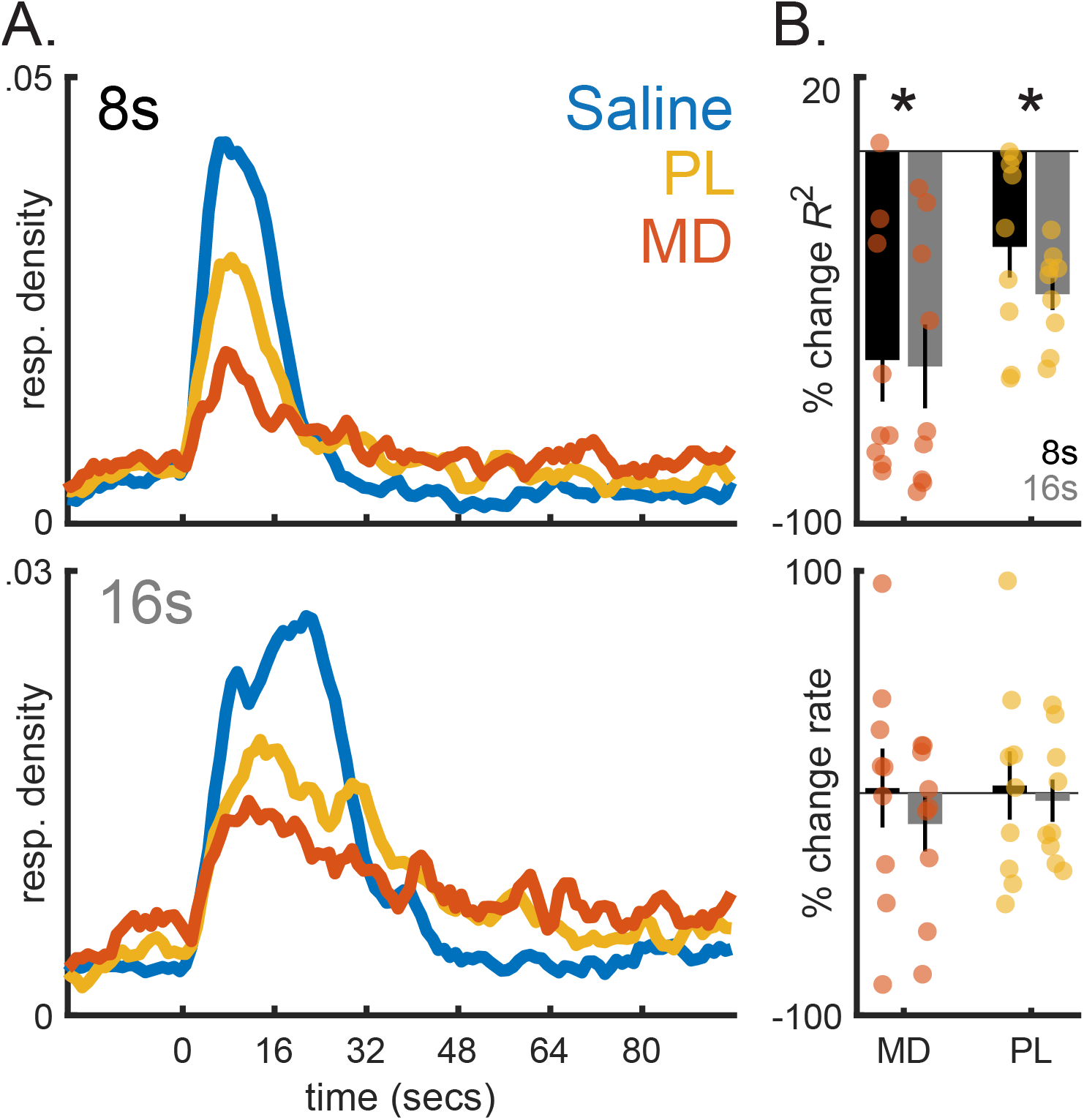
Inactivating the prelimbic cortex (PL) or mediodorsal thalamus (MD) flattens mean response curves (*n* = 9). (A) Mean response rate across time during probe trials (normalized by sum) for the 8s and 16s cues (top and bottom, respectively), split by infusion type. Data were grouped into 1s bins and have been smoothed over a 3-bin window (for visualization only). (B) Percent change in the fit-quality of a gaussian distribution to response rates across time and overall mean response rates (i.e., ignoring time) during probe trials for each area, relative to saline sessions (top and bottom, respectively). Asterisks indicate *p* < 0.05, relative to saline sessions for a given area.

While these data show a clear timing deficit, they do not necessarily imply that the MD and PL are required for organizing behavior in time. The reason relates to the fact that, during single-trials in this task, rats’ responses do not show a gaussian shape, unlike the averaged response curves above. Rather, rats emit a discrete ‘burst’ of responses around the entrained interval, with a sharp onset/offset (Church et al., 1994). When this burst starts and stops usually varies slightly from trial to trial. Consequently, when averaged across trials, mean response rates show a smooth, Gaussian shape (see Fig 1A for visual illustration). Flatter mean response curves might imply that the burst-pattern was also disrupted. However, if our inactivations increased the variability of when the burst occurred, response curves would also appear flat.

To dissociate these possibilities, we analyzed single-trial responding next. Figure 3A shows heat maps of responses across time for all probe trials collected in the study. We quantified the burst using the standard approach, wherein a step function is fit to single-trial responses (see Methods; Church et al., 1994). Importantly, even if response times are entirely random, bursts will occasionally form by chance. The question is whether the bursts are ‘coincidental’ or a meaningful component of the data. For illustration, we also ran the analysis on time-shuffled data from saline sessions and present a heat map.

**Figure 3.**
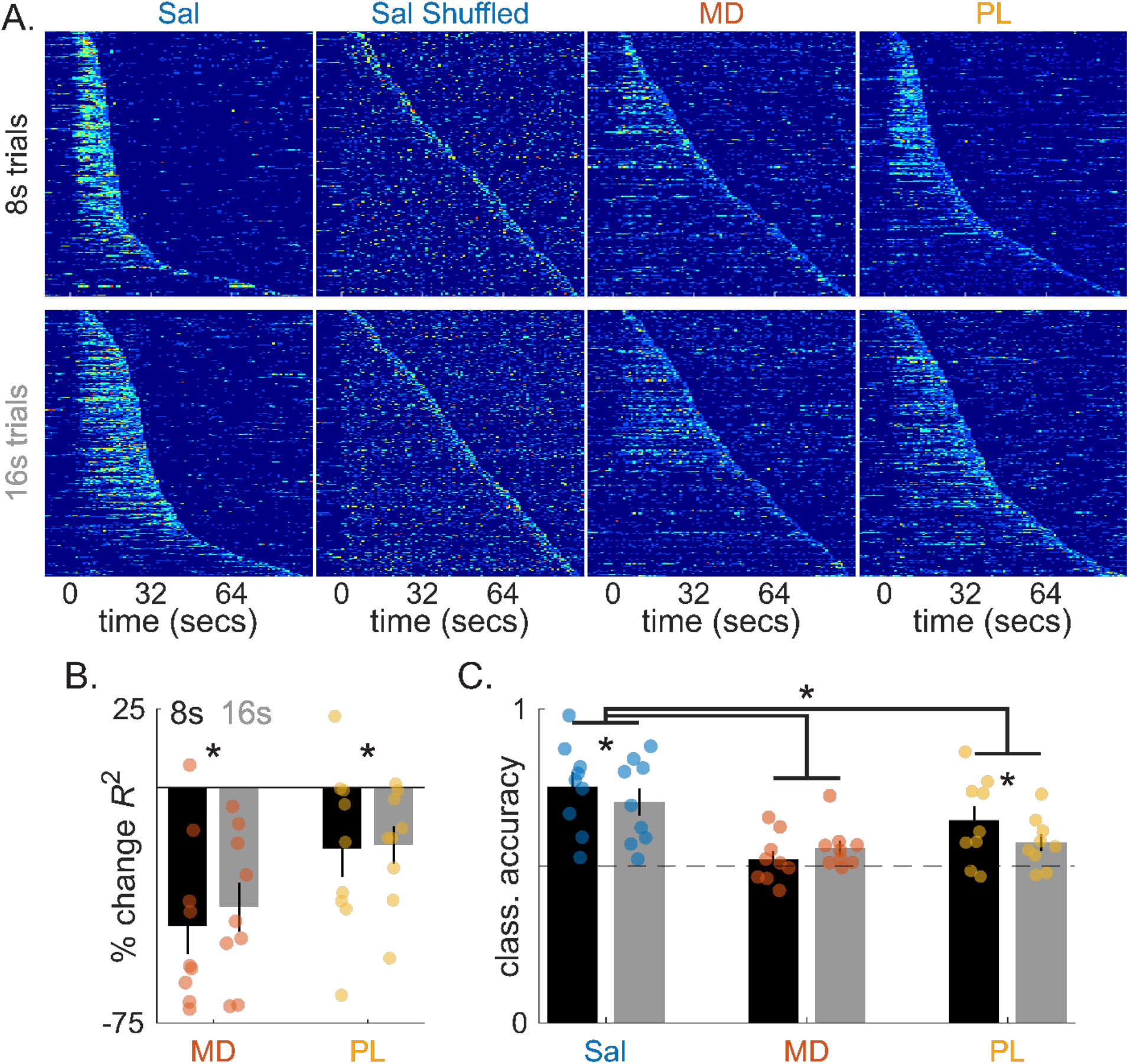
Inactivating the MD or PL disorganizes singe-trial patterns of timed responding (*n* = 9). (A) Heat maps of responses during individual trials for each infusion type (1s bins). Shuffled saline data is also presented (see text). All data have been sorted by the stop-time from the burst analysis. (B) Percent change in fit-quality of a step function fit to individual trials for each inactivation type, relative to saline. (C) Accuracy of a support vector machine trained to discriminate shuffled data from unshuffled data, split by infusion type. Centered asterisks indicate above-chance classification for a given infusion type (dotted-line = .5). Asterisks above horizontal lines indicate significance across areas.

Data from saline sessions show a sharp distinction between the pre/post burst periods (Fig 3A). The contrast is less clear in the shuffled data, as bursts were detected randomly across time. Importantly, the burst pattern is also less apparent in the MD and PL inactivation data, with rats showing a lower density of responses within the burst period, along with an elevation in sparse responses/additional bursts throughout the trial. Consistent with this, the fit-quality of a step-function to individual trials dropped following both MD and PL inactivations [Fig 3B; Infusion: *F*(2,14) = 13.34, *p* < 0.005; MD: *t*(6) = -4.38, *p* < 0.005; PL: *t*(6) = -2.51, *p* < 0.05; *R*^2^ *M* +/-*SEM*: Sal = 0.72 +/-0.03, MD = 0.46 +/-0.03, PL = 0.61 +/-0.03]. As a more direct test, we trained a machine learning algorithm (support vector machine) to discriminate shuffled and unshuffled single-trial data for each infusion type. This provides a ‘data driven’ approach to evaluating the organization of behavior, casting few prior assumptions regarding how the data should be organized, per se. As expected, accuracy was high for saline data and dropped following MD and PL inactivations [Fig 3C; Infusion: *F*(2,14) = 10.16, *p* < 0.005; MD: *t*(6) = -3.77, *p* < 0.01; PL: *t*(6) = 3.22, *p* < 0.05; *M* +/-*SEM*: Sal = 0.73 +/-0.03, MD = 0.54 +/-0.02, PL = 0.61 +/-0.03]. While accuracy under MD and PL inactivations began to approach chance, all infusion types were either significantly above chance, or marginally so in the case of the MD [Sal: t(8) = 7.09, p < 0.001, MD: t(8) = 2.12, p = 0.067, PL: t(8) = 3.65, p < 0.01]. As a whole, these data indicate that the MD and PL are necessary for guiding behavior based on time-information.

### Communication between the MD and PL is necessary for timing performance

The MD and PL share dense reciprocal connections and often co-project to areas known to be involved in timing (e.g., striatum). The above data clearly show that both areas play a role in timing. However, whether they act independently or interact during timing tasks is still an open question. Therefore, in our next experiment, we tested whether MD-PL communication, specifically, is involved in timing.

Our general approach was identical to the prior experiment. We trained 10 rats on the same task and implanted cannulae bilaterally in the MD and PL. However, instead of inactivating either area bilaterally, we ran a ‘disconnection’ procedure (Fig 4A; Barker et al., 2007). Specifically, all muscimol inactivations targeted both areas unilaterally. Importantly, our inactivations were either ipsilateral (e.g., right-PL + right-MD) or ‘crossed’ (e.g., left-PL + right-MD). Ipsilateral inactivations should have weak effects, as the two areas can still communicate in the contralateral hemisphere. By contrast, crossed inactivations will disrupt MD-PL communication bilaterally. Therefore, if MD-PL communication mediates performance, crossed infusions should yield a large deficit. Notably, in both cases, any independent communication that the MD or PL makes with other areas will be preserved unilaterally. Therefore, this design provides a selective means of disrupting mono-or poly-synaptic MD-PL interactions.

**Figure 4.**
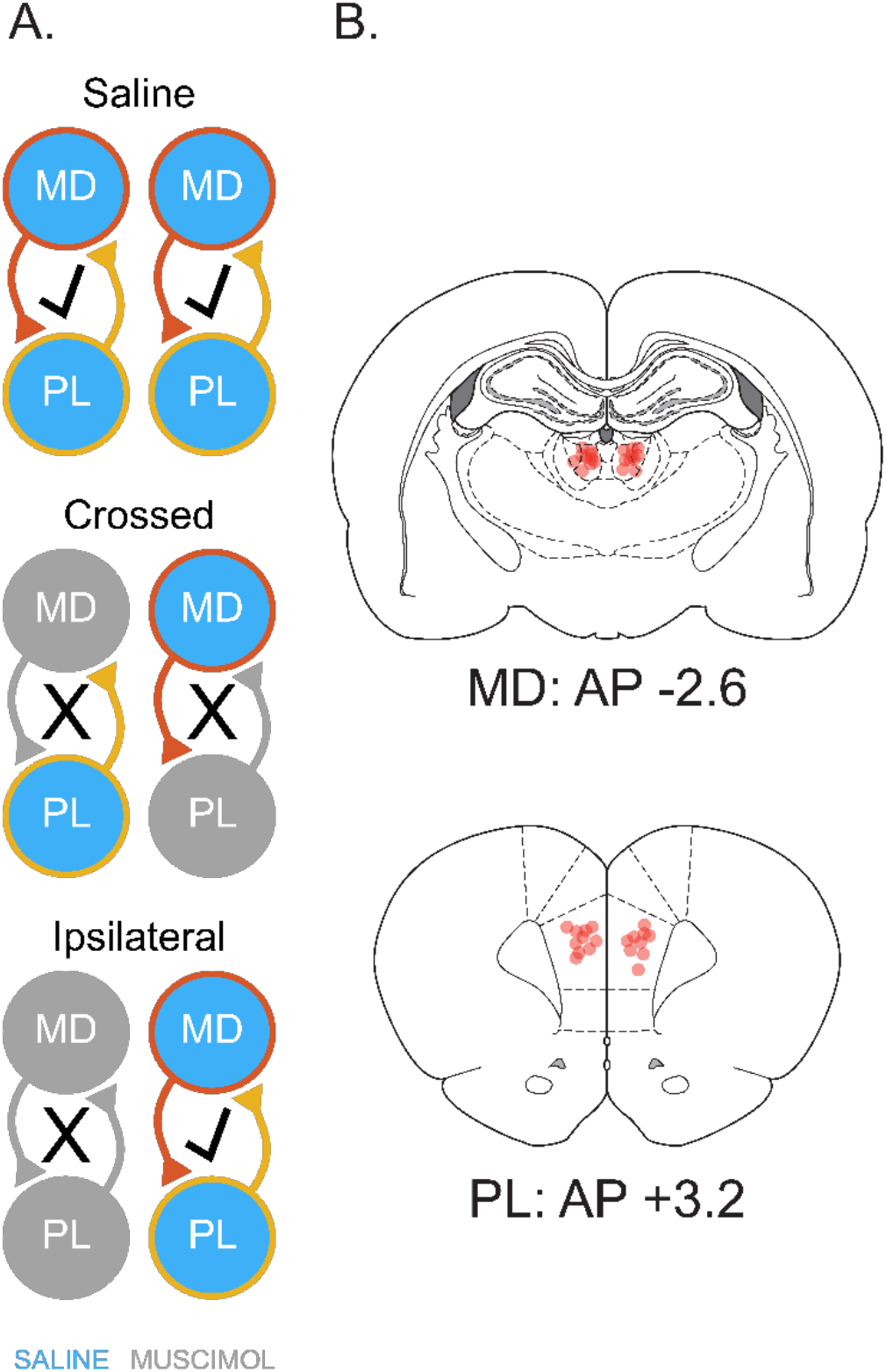
Inactivation design and histological analysis for MD-PL disconnection procedure. (A) Illustration of infusion design for the disconnection procedure, including saline, crossed, and ipsilateral infusions. For simplicity, only one spatial permutation for the crossed and ipsilateral infusions is diagramed. However, all permutations were performed across sessions (see methods). (B) Diagram of infusion cannula tip locations for all rats in the experiment for the MD (top) and PL (bottom).

During saline sessions, mean probe-trial response rates showed the typical gaussian shape for both trial types (Fig 5B). Furthermore, ipsilateral inactivations had no significant effects on performance. Importantly, crossed infusions flatten the curves, similar to bilateral inactivations in the prior experiment. Consistent with these impressions, we observed a drop in Gaussian fit-quality during crossed infusion sessions alone, relative to saline and ipsilateral inactivations [Fig 5B; Infusion: *F*(2,16) = 15.53, *p* < 0.005; Sal vs. Cross: *t*(7) = -3.57, *p* < 0.05; Cross vs. Ipsi: *t*(7) = -4.42, *p* < .005; Sal vs. Ipsi: *t*(7) = 2.22, *p* = .057; *R*^2^ *M* +/-*SEM*: Sal = 0.94 +/-0.13, Cross = 0.71 +/-0.13, Uni = 0.95 +/-0.13]. Furthermore, inactivations did not reliably impact overall trial response rates [Infusion: *F*(2,16) = 0.613, *p* = 0.487].

**Figure 5.**
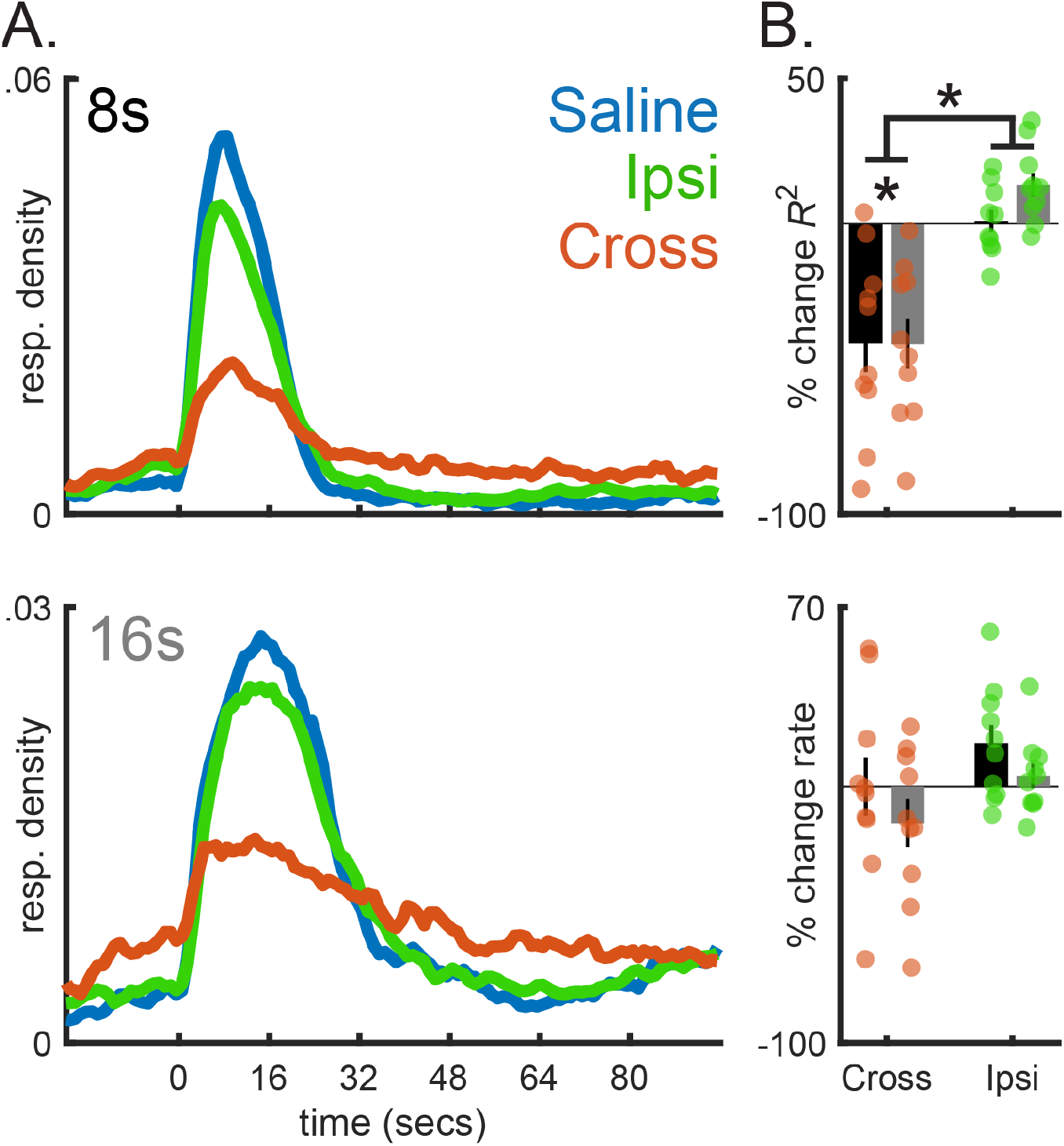
Disconnecting the MD from the PL flattens mean response curves (*n* = 10). (A) Mean response rate across time during probe trials (normalized by sum) for the 8s and 16s cues (top and bottom, respectively), split by infusion type. Data were grouped into 1s bins and have been smoothed over a 3-bin window (for visualization only). (B) Percent change in the fit-quality of a gaussian distribution to response rates across time and overall mean response rates (i.e., ignoring time) during probe trials for each inactivation type, relative to saline sessions (top and bottom, respectively).

Similar to individual inactivations of the MD or PL, blocking MD-PL communication disrupted the burst pattern of responding during single-trials (Fig 6A). Specifically, crossed infusions selectively dropped the fit-quality of a step function to individual trials [Fig 6B; Infusion: *F*(2,16) = 24.15, *p* < 0.001; Sal vs. Cross: *t*(7) = -4.75, *p* < 0.005; Ipsi vs. Cross: *t*(7) = -6.65, *p* < .001; Sal vs. Ipsi: *t*(7) = -1.11, *p* = .300; *R*^2^ *M* +/-*SEM*: Sal = 0.75 +/-0.02, Cross = 0.53 +/-0.03, Ipsi = 0.73 +/-0.01]. Regardless of infusion-type, overall fit quality was higher for the 8s cue [Cue: *F*(1,8) = 69.48, *p* < 0.001]. However, crossed infusions attenuated this difference, likely related to a floor-effect [Cue X Infusion: *F*(2,16) = 6.38, *p* < 0.01; 8s vs. 16s cue: Sal: *t*(8) = 8.82, *p* < 0.001; Cross: *t*(8) = 2.30, *p* = 0.05; Ipsi: *t*(8) = 8.15, *p* < 0.001]. We found similar results when training a machine learning algorithm to discriminate shuffled from unshuffled data. Specifically, crossed inactivations decreased accuracy, relative to both saline and ipsilateral infusions [Fig 6C; Infusion: *F*(2,16) = 47.70, *p* < 0.001; Sal vs. Cross: *t*(7) = -8.29, *p* < 0.001; Ipsi vs. Cross: *t*(7) = 7.88, *p* < 0.001; Sal vs. Ipsi: *t*(7) = 2.30, *p* = 0.05; *M* +/-*SEM*: Sal = 0.79 +/-0.03, Cross = 0.58 +/-0.02, Ipsi = 0.74 +/-0.02]. In all cases, overall accuracy was above chance [Fig 6C; Sal: *t*(9) = 12.33, *p* < 0.001, Cross: *t*(9) = 5.06, *p* < 0.005, Ipsi: *t*(9) = 14.07, *p* < 0.001]. However, we observed better overall accuracy for the 8s cue with crossed inactivations showing a weaker difference, again suggesting a floor effect [Fig 4c; Cue X Infusion: *F*(2,16) = 8.90, *p* < 0.005; 8s vs 16s: Sal: t(8) = 7.53, p < 0.001; Cross: t(8) = 3.08, p < 0.05; Ipsi: t(8) = 5.29, p < 0.005].

**Figure 6.**
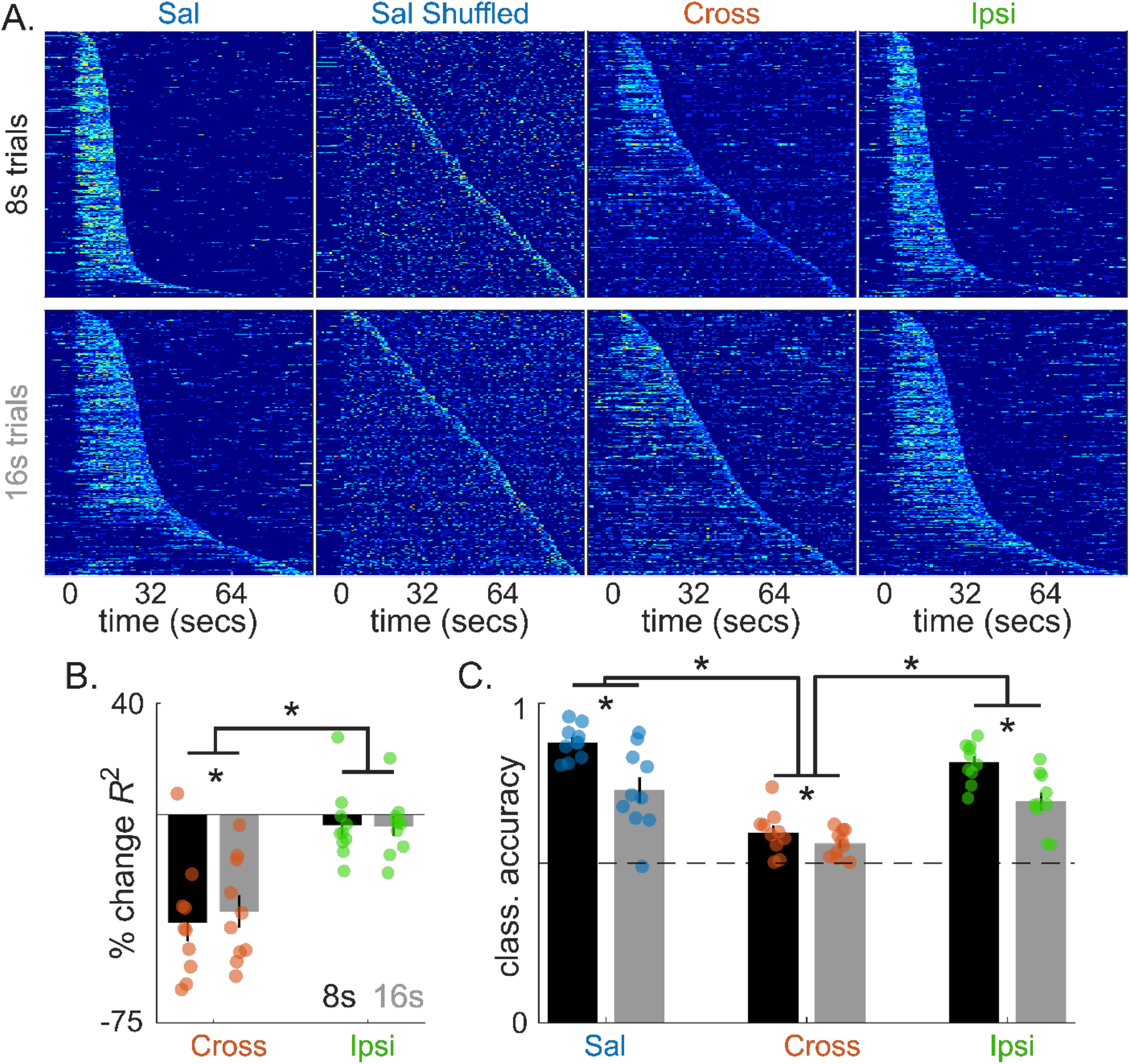
Disconnecting the MD from the PL disorganizes singe-trial patterns of timed responding (*n* = 10). (A) Heat maps of responses during individual trials for each infusion type (1s bins). Shuffled saline data is also presented (see text). All data have been sorted by the stop-time from the burst analysis. (B) Percent change in fit-quality of a step function fit to individual trials for each inactivation type, relative to saline. (C) Accuracy of a support vector machine trained to discriminate shuffled data from unshuffled data, split by infusion type.

As a whole, these data confirm that communication between the MD and PL mediates timing behavior.

## Discussion

Our data suggest that the mediodorsal thalamus (MD), prelimbic cortex (PL), and MD-PL interactions are necessary for timing performance. Inactivating either area or blocking MD-PL communication prevented subjects from effectively organizing their behavior in time, with no apparent effect on motivation or ability to respond, per se. These data extend prior work suggesting that the MD and PFC interact to regulate adaptive behavior. For example, MD-related cognitive deficits often resemble PFC dysfunction, such as impaired executive function, working memory, etc. (Golden et al., 2016; Parnaudeau et al., 2018). Furthermore, MD inactivation disrupts PFC activity during cognitive tasks and vice versa (Alexander & Fuster, 1973; Parnaudeau et al., 2013). However, data supporting this hypothesis have been primarily correlative, monitoring changes in purported task-related activity in one area after inactivating the other. The data presented here contributes to the recent, but growing, body of causal data showing that MD-PFC interactions mediate cognitive function (Bolkan et al., 2017; de Kloet et al., 2021).

Our data may be particularly valuable for studying the MD using rodents. Specifically, establishing a reliable rodent task for evaluating the MD’s role in cognition has been challenging. Prior work has primarily focused on spatial working-memory tasks. Several studies have indeed found effects of MD manipulations on performance (Bailey & Mair, 2005; Bolkan et al., 2017; Young et al., 1996). However, the literature is admittedly ‘mixed,’ as many others have reported null findings (Burk & Mair, 1998; Hunt & Aggleton, 1998a, 1998b; Neave et al., 1993). The peak-interval procedure may be a promising alternative assay, yielding large effects for both MD/PL inactivations and MD-PL disconnection. The task also recruits similar cognitive sub-processes to spatial tasks, such as working memory (Buhusi & Meck, 2006), attention (Buhusi & Matthews, 2014), and decision-making (MacDonald et al., 2012). Furthermore, as discussed below, deficits have been reliable across studies thus far.

This could have important implications for public health research. For example, schizophrenia produces timing deficits (Densen, 1977; Lee et al., 2009; Singh et al., 2019). Impaired MD-PFC communication is already implicated in more general cognitive impairments seen in this disorder (Giraldo-Chica et al., 2018; Parnaudeau et al., 2018). Therefore, a natural question is whether the two deficits are related. The peak-interval procedure provides an added advantage here, as the task translates easily between rodents and humans with minimal modifications needed (Malapani et al., 2002; Rakitin et al., 1998). Therefore, future work could explore this topic in humans using fMRI and/or EEG, and more invasive techniques can be employed with rodents.

These data also give further insight into the networks that support timing, which are currently unclear. With respect to prior work, Meck (2006) observed a behavioral effect that closely parallels ours after lesioning midbrain dopamine centers during the peak-interval task with rats. Specifically, much like our data, rats maintained their response rates during the cue-period, yet were unable to control response timing, flattening the curves.

Meck (2006) hypothesized that dopamine signaling functions as a ‘start-gun,’ activating downstream timing circuitry when cues are presented. When this signal is blocked, the timing circuitry fails to activate at cue-onset. Therefore, responses are purely guided by motivational systems, which are generally independent of timing (Kurti & Matell, 2011; Roberts, 1981).

These data might extend Meck’s (2006) results. For example, if the MD and PL are truly critical timing nodes, our manipulations would constitute direct inactivation of the timing network itself. As a slight alternative, both the MD and PL are densely innervated by midbrain dopamine centers (Descarries et al., 1987; García-Cabezas et al., 2007).

Therefore, there could be a more direct link between Meck’s lesions and our data. These data also extend prior work studying the MD or PL in timing individually. For example, Buhusi et al. (2018) inactivated the PL in a dual, visual-cue peak task, similar to the one used here. Importantly, they used graded doses of muscimol, ranging from low to moderate (.11 and 1mM, respectively), to evaluate progressive PL dysfunction on timing. As the dose increased, the spread of the response curves increased. As we were interested in necessity, we aimed for asymptotic inactivations with a 2mM muscimol dose. This is still considered moderate with respect to the broader literature, with ‘high’ being in the 5-8mM range (Katz et al., 2016; Randall et al., 2021). However, our pilot data suggest that higher doses do not increase effect sizes. We avoided these doses over concerns that, during MD infusions, small leaks might enter the ventrolateral thalamus. In our experience, VL inactivation produces profound dystonia in rats (see also Di Chiara et al., 1979).

It would be convenient to argue that our data represent a simple extension of Buhusi et al.’s (2018) spread effect, with the spread broadening further across the trial period. However, this would imply that the burst-pattern was preserved during single-trials in our data. Visual inspection of the heat maps certainly suggests that bursting occasionally occurred. However, our analyses suggest that the primary component of MD/PL disruption is a breakdown in the organization of timing behavior during single-trials. Nonetheless, a reasonable assumption is that, when the timing system becomes sufficiently impaired, behavioral control is transferred to motivational systems, as discussed above.

Another relevant study comes from Lusk et al. (2020). Using mice, they performed MD muscimol inactivations during the peak procedure with a single visual-cue/interval. Due to either a species difference or a smaller dose (.08mM), they found adequate preservation of Gaussian-like response rates. Similar to Buhusi et al. (2018), they observed an increase in spread. However, they also saw an increase in peak times. This could suggest that the MD and PL play partially independent roles in timing. Indeed, in our data, effect sizes appear larger following MD inactivations or MD-PL disconnections, relative to the PL inactivations alone. However, we urge caution with such comparisons, as the MD is physically smaller than the PL. Therefore, a plausible alternative is that we were able to achieve greater coverage of the MD, leading to larger effects. Nonetheless, if accepted, these data could indicate that the MD interacts with other cortical areas that often receive less attention in the rodent timing literature (e.g., cingulate, premotor, etc.).

This study opens several further avenues for future research. For example, an important direction is to more clearly define how the MD and PL interact during timing tasks. A straightforward proposal is that, when task-related cues are presented, the MD relays this information to the PL, activating timing circuitry it contains. Consistent with this, during working memory tasks, there is some indication that the MD initiates maintenance circuitry in the PFC, whereas the PFC mediates later actions based on the maintained information (Bolkan et al., 2017). This would also account for our data, as one would expect MD/PL inactivation or disconnection to produce a comparable deficit. Furthermore, the MD would still serve the thalamus’ known role as a sensory relay, yet selectively processing learned cues related to adaptive behavior. However, feedback projections from the PFC to the MD appear to mediate cognitive performance (Bolkan et al., 2017), and the MD may very well play a more active, continuous role than this view would imply. Furthermore, poly-synaptic feedback loops from the PFC to the striatum could also play a role, as the basal ganglia innervate the MD (Georgescu et al., 2020). Disconnection designs do not provide sufficient selectivity to dissociate these possibilities. Therefore, projection targeting with optogenetics or DREADDs would be very useful in this regard.

Finally, while strong emphasis is not given to this point in the current manuscript, the present task incorporated cross-modal cues (i.e., tone and light). We did not find preferential inactivation effects on visual or auditory stimuli, although there were occasional baseline modality-duration interactions that acted independently of infusions (see supplemental results). This suggests that the PL and MD play amodal roles in timing. This extends a lasting question over whether timing is processed by a centralized, amodal network or by distributed, sensory-specific systems (Paton & Buonomano, 2018)—favoring the former. Interestingly, time-related signals have been documented in primary sensory areas (e.g., primary visual cortex; Shuler, 2006). Therefore, these data also open the question as to how information from distinct modalities converges onto the timing network characterized here.

## Methods

### Behavioral training

All training protocols follow closely from De Corte et al. (2018), and were consistent with the University of Iowa’s Animal Care and Use Committee guidelines and the Declaration of Helsinki. We trained male Long-Evans rats (total *n* = 19) in standard operant chambers (MedAssociates) under modest water deprivation. Body weights were monitored and maintained to 90% of non-deprived levels. Training progressed through three phases. During the initial ‘shaping’ phase (3 sessions), rats learned to make responses on a nose poke to earn water reward (0.057 ml). Sessions lasted 60 minutes (mins). No cues were included in this phase, and any response on the nose poke yielded reward. Next, a ‘fixed-interval’ training phase began (6 sessions; 120 mins each). During this phase, trials began with the presentation of either a tone (4.5 kHz; 80 dB) or houselight. Each cue predicted reward availability after either an 8 or 16 second (s) duration elapsed (Fig 1A). We counterbalanced the modality-duration relationship across rats (e.g., tone-8s / light-16s vs. light-8s / tone-16s). Once the rat made a response after the respective cue’s interval passed, the cue terminated and reward was delivered. Early responses were not penalized. Trials were followed by a dark inter-trial interval, lasting 90s on average (30s minimum + a variable, 60s geometrically distributed interval). After fixed-interval training, a ‘probe training’ phase began. Probe training was identical to fixed-interval training. However, during a subset of ‘probe trials,’ the cue remained on for 96-128s, regardless of when rats responded, and no reward was delivered (Fig 1A). Probe trials were randomly intermixed with rewarded trials. We gradually increased the proportion of probe trials during this phase (25% for the first 3 sessions and 50% for the remaining 6 sessions). Furthermore, the ITI was gradually decreased to 45s (30s-fixed + 15s-geometric). After initial training, rats underwent surgery.

### Surgery/Histological analysis

Rats received surgery under isoflurane anesthesia (5% induction; 2.5% maintenance). We implanted four 23-gauge guide cannulae bilaterally over either the prelimbic cortex (PL) or mediodorsal nucleus (MD) of the thalamus (PL: AP: +3.2, ML: +/-0.7, DV: -2.0; MD: AP: -2.6 ML: +/-0.7 DV: -4.4; relative to bregma/skull surface). We also inserted 30-gauge dummy cannulae that extend 1mm beyond the base of the guide cannulae. To shield the junction between the guide’s top and the dummy cannulae, we placed 19-gauge tubing on the outside of the guides, extending 3mm above the top. Infusion cannulae used for drug delivery (see below) extended 1.5mm beyond the base of the guide. All cannulae were stainless steel 304 and custom-fabricated (Component Supply Inc.). We secured the implants with dental cement, and rats recovered in their home cage for 1 week without water deprivation before further experimentation. After the experiment, we perfused rats using chilled saline followed by 4% paraformaldehyde (120ml for both). Brains were extracted, stained with thionin, and imaged under a microscope to verify cannula placement.

### Retraining and restraint habituation

Following recovery, we retrained rats on the task for 7 sessions. During this time, we gradually acclimated them to restraint procedures used for drug infusions (De Corte et al., 2019). Specifically, on day 1, we placed rats in a .5’ X 1’ plexiglass enclosure, where all infusions took place, and allowed them to move freely for 1 minute. On day 2, we lightly restrained the rats by hand in the enclosure for 1 minute. On days 3 and 4 the restraint time increased from 3 to 5 minutes, respectively. On days 5 and 6, we conducted mock-infusions that paralleled procedures used during actual infusion sessions (see below). Specifically, we removed dummy cannulae and cleaned them with 70% ethanol. Then, we inserted two 30-gauge infuser cannulae into the guides (one PL and one MD; 1.5mm extension from the guide-base). We ran an infusion pump (Stoelting), connected to the infusers by polyethylene tubing, at a rate of .1 ul/min for 2 minutes, although no substance was loaded in the infusers. Once the pump finished, we left the infusers in place for 1 minute. Subsequently, we switched the infusers to the remaining 2 PL/MD guide cannulae and repeated the procedure. No restraint habituation was conducted on the 7th day because all actual infusion sessions were preceded by normal behavioral recovery days.

### Drug infusions

Following retraining and restraint habituation, infusion sessions began. All protocols were identical to the mock infusion sessions described above. However, we now infused .22ul of either saline or 2mM muscimol (a GABA-A receptor agonist; Sigma Aldrich). Our infusion targets differed by experiment. Specifically, for the bilateral inactivation experiment, we targeted the PL or MD bilaterally with muscimol and infused saline into the other brain area. For baseline sessions, we infused saline at all targets. This yielded three infusion types (muscimol-PL / saline-MD; saline-PL / muscimol-MD; saline-PL / saline-MD). For the disconnection experiment, saline sessions still targeted all four sites. However, during muscimol sessions, we always targeted one PL and one MD for inactivation. Specifically, during ‘ipsilateral’ inactivation sessions, the inactivated PL and MD were in the same hemisphere (left-PL / left-MD; right-PL / right-MD). Conversely, during ‘crossed’ inactivation sessions, the inactivated PL and MD were in opposing hemispheres (left-PL / right-MD; right-PL / left-MD). In both cases, we infused saline into the remaining areas. Rats began the task 30 mins after the final infusion took place. All rats received each ipsilateral/crossed infusion combination in a randomized order. One session of normal training separated all infusion sessions to allow for behavioral recovery and prevent GABA receptor sensitization.

### Behavioral measures

#### 1) Mean response rate analysis

We analyzed mean response rates by grouping responses during probe trials into 1s bins and averaging across trials. To quantify timing behavior, we fit each response curve with a Gaussian + kurtosis parameter function described in the following equation (De Corte et al., 2018; Swanton & Matell, 2011):

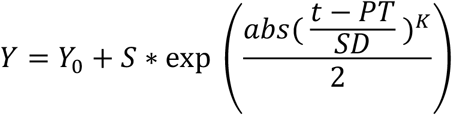

*Y*_*0*_ represents the baseline rate of responding, *S* is a scale parameter, *t* is time, *PT* is the mean (i.e., ‘peak-time’), *SP* is the spread, and *K* allows the function to fit peaks with varying degrees of kurtosis/skew. As described below, we were primarily interested in the fit-quality, measured by the *R*^2^. Note that, for all statistical analyses on *R*^2^ data, we used the Fisher’s *R* to *Z* conversion on the square root of each value. This is standard and required to obtain normality in data based on Pearson’s correlation coefficients. We also report peak-time and coefficient of variation (spread / mean) measures from the fits in the supplemental results section. Finally, to assess changes in motivation or motor function during inactivation sessions, we also computed the mean response rate during the trial-period (i.e., the mean, ignoring time).

#### 2) Single-trial analyses

We analyzed single-trial patterns of responding, again using 1s bins. We identified purported response bursts using a standard approach in which a dual step-function (first step-up, second step-down) is fit to responding (Church et al., 1994). The algorithm finds bursts of responding by iteratively fitting the step function across time in the trial, minimizing absolute residuals. Our measures of interest were the fit-quality of the function (*R*^2^), as well as the start and stop times, defined as the time where the initial up-step and the terminal down-step were found, respectively. For the machine learning analysis, we trained a support vector machine (SVM) to discriminate the original single-trial data from the same data yet time-shuffled (i.e., binary classification; Matlab, fitcsvm function). The primary feature was time (i.e., bin 1s, 2s, etc.), and each trial served as a set of observations. For binary classification, SVMs segregate the two data sets with a non-linear hyperplane when a kernel function is used. We used a standard, radial basis function to compute the boundary. For each rat, we trained the SVM using the original and shuffled data, aside from one trial, and then tested the classifier with the unused trial (i.e., leave-one-out cross-validation). The analysis results will differ slightly by chance, due to random shuffling. Therefore, to obtain a more robust estimate of mean accuracy, we repeated each analysis 30 times—the average number of probe trials for each cue, per infusion type—and took the mean accuracy across all runs. However, incorporating more/less runs did not change the results.

### Statistical Analyses

We analyzed each measure using a mixed-model Analysis of Variance (SPSS). We assessed sphericity using Mauchly’s test. In cases where sphericity was violated, we employed the Greenhouse-Geisser correction. For both experiments, modality-duration pairing (tone-8s / light-16s vs. light-8s / tone-16s) served as a between-subjects factor. For the bilateral inactivation experiment, Infusion (saline, MD muscimol, PL muscimol) and Cue (8s vs. 16s) served as within-subjects variables. The model was the same for the disconnection experiment. However, the Infusion factor corresponded to the spatial arrangement of the inactivations (saline, ipsilateral, crossed). To simplify reporting, we assessed laterality effects with individual ANOVAs (e.g., Crossed: left-MD / right-PL vs. right-MD / left-PL). As expected, there were no reliable effects [all *F*s < 1.75]. Therefore, we pooled the data. We used simple-effects analyses to probe main effects or interactions and report the corresponding *t* values/degrees of freedom. Where appropriate, we used Dunnet’s test to control for multiple comparisons.

