## supplemental data for "Communication between the mediodorsal thalamus and prelimbic cortex regulates timing performance in rats"

**Supplemental results.**

*Traditional timing measure analyses.*

Across both experiments, we note a marked decrease in fit-quality following inactivations of the MD and PL or after disconnecting the two areas. This held for both mean response rate analysis (i.e., Gaussian fits) and the single trial analyses (i.e., step-function fits). Moreover, the data overall suggest that the typical patterns of responding, on which these analyses were developed, broke down following inactivations. Consequently, it is difficult to interpret the meaning of traditional timing measures derived from these analyses, and they will, of course, be outlier-prone. Nonetheless, we report standard results here for exploratory purposes.

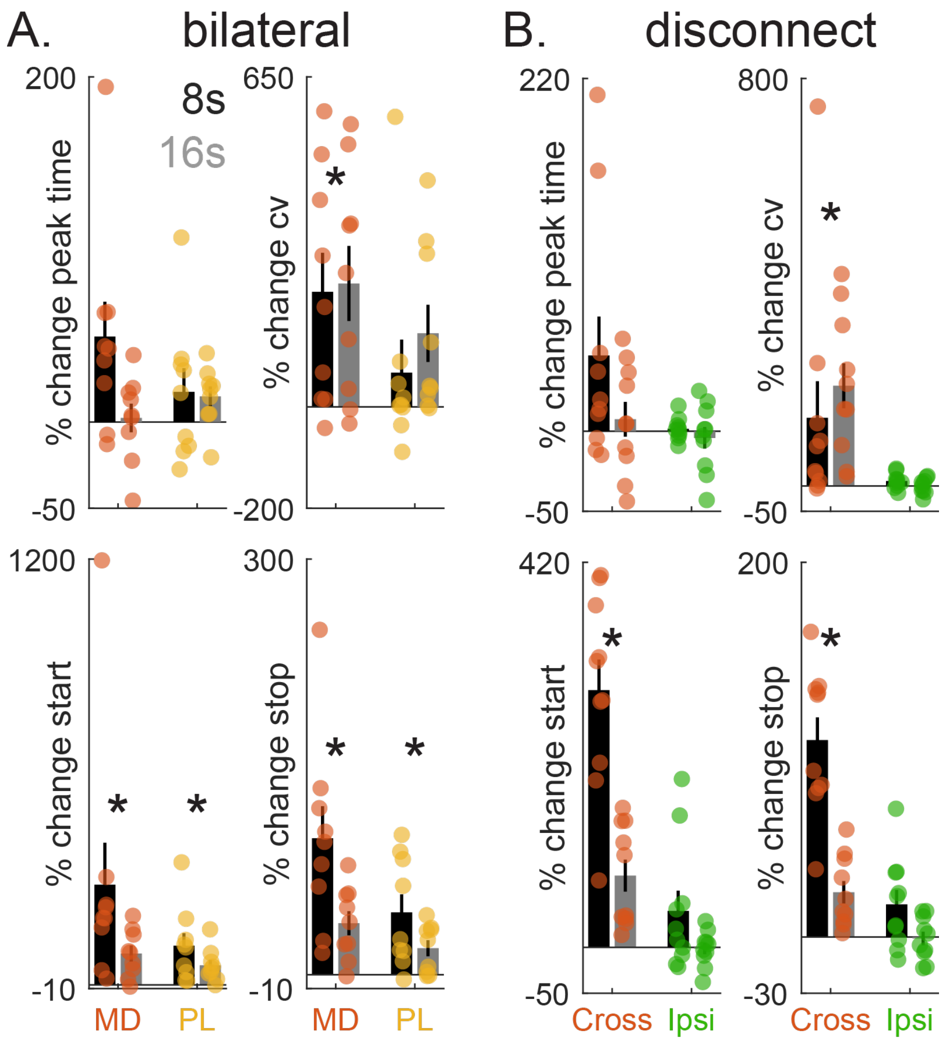


***Figure 7.*** **Impacts of MD/PL inactivation or disconnection on traditional timing measures.** All data are presented as percent change, relative to saline sessions. (A) Data from the bilateral inactivation experiment. Peak time and CVs (i.e., spread over mean) from the Gaussian fits to mean response rates are presented at the top. Single trial start- and stop-times from the burst analysis are presented at the bottom. (B) same as (A) but data come from the disconnection experiment.

For the bilateral inactivation experiment, we did not observe significant peak-time effects or interactions related to inactivations [Fig 2d; Fs < 2.5, ps < 0.05]. Inactivations did elevate overall CVs (i.e., spread/peak-time), although this was only significant for the MD [Fig 7A2e; Infusion: $F(2,14) = 7.08$, $p < 0.01$; MD: $t(6) = 3.69$, $p < 0.01$; PL: $t(6) = 1.78$, $p = 0.117$]. With respect to single-trial analyses, we observed an increase in start and stop times [Fig 7A(bottom)2d; Start: Infusion: $F(2,14) = 17.14$, $p < 0.001$; MD: $t(6) = 5.13$, $p < 0.005$; PL: $t(6) = 5.56$, $p < 0.005$; Stop: Infusion: $F(2,14) = 19.01$, $p < 0.001$; MD: $t(6) = 5.70$, $p < 0.005$; PL: $t(6) = 4.22$, $p < 0.005$]. Note that this pattern is generally consistent with more uniform/random detection of start and stop times across the trial period, as all measures will pull toward the center of the trial (i.e., 48s). Examples can be seen in the shuffled saline data in Figures 3a and 5a.

For the disconnection experiment, we found no significant peak time changes related to infusions [*F*s < 2 ; *p*s > 0.05]. Furthermore, crossed inactivations elevated CVs [Infusion: $F(1,8) = 9.29$, $p < .05$; Sal vs. Crossed: $t(7) = 3.03$, $p < .05$; Ipsi vs. Crossed: $t(7) = 3.07$, $p < .05$; Sal vs. Ipsi: $t(7) = -0.14$, $p = .895$]. With respect to single-trial analyses, we found a main effect of infusions on start times that interacted with cue [Fig 7B; Infusion: $F(2,16) = 51.01$, $p < 0.001$; Cue X Infusion: $F(2,16) = 12.26$, $p < 0.005$]. This resulted from start times being later for the 16s cue during saline and ipsilateral infusions, as expected, but not reliably different for crossed infusions [Sal: $t(8) = 9.14$, $p < 0.001$; Cross: $t(8) = 1.53$, $p = 0.164$; Ipsi: $t(8) = 5.88$, $p < 0.001$]. The same pattern was found for stop times, although the cue difference was now marginally significant following crossed inactivations [Infusion: $F(2,16) = 46.46$, $p < 0.001$; Cue X Infusion: $F(2,16) = 16.40$, $p < 0.001$; 8s vs. 16s cue: Sal: $t(8) = 12.34$, $p < 0.001$; Cross: $t(8) = 2.01$, $p = 0.08$; Ipsi: $t(8) = 8.87$, $p < 0.001$]. Similar to the mean-effects seen in the bilateral experiment, these data are broadly consistent with more uniform/random start-time detection across the trial for both cues during crossed infusions (i.e., 'pulling' toward the center timepoint of the trial).

*Incidental ANOVA findings.*

In the primary results section, we focus on effects or interactions related to inactivations. However, there were cases in which the omnibus ANOVA revealed incidental findings that were unrelated to infusions. These effects were generally sporadic (i.e., did not replicate across experiments) and secondary to the focus of this manuscript. Nonetheless, we report them here for those who may be interested.

First, for the bilateral inactivation experiment, the tone-8s / light-16s group responded slightly more to the 16s cue than the light-8s / tone-16s group. No differences were found for the 8s cue, yielding a Cue X Modality-Duration Pairing interaction [Cue X Modality-Duration Pairing: $F(2,14) = 8.19$, $p < 0.05$; Short: $t(7) = 0.470$, $p = 0.653$; Long: $t(7) = -3.37$, $p < 0.05$]. We found no such differences in the disconnection experiment.

Second, in the disconnection experiment, rats in the tone-8s / light-16s group showed a higher Gaussian fit quality for the long cue than the tone-16s / light-8s group, yielding an interaction [Cue: $F(1,8) = 38.41$, $p < 0.005$; Cue X Modality-Duration: $F(1,8) = 15.89$, $p < 0.005$; Short: $t(8) = 0.51$, $p = 0.621$; Long: $t(8) = -2.37$, $p < 0.05$]. Correspondingly, when training the machine learning algorithm to discriminate shuffled from unshuffled single-trial data, the tone-8s / light-16s group showed a higher decoding accuracy for the long cue than the tone-16s / light-8s group [Cue X Modality Duration-Pairing: $F(1,8) = 27.84$, $p < 0.005$; 8s cue: $t(8) = 1.02$, $p = 0.338$; 16s cue: $t(8) = -2.78$, $p < 0.05$]. No modality group effects were found throughout the bilateral inactivation experiment analysis.
